# Gut microbiome species *Levilactobacillus brevis* regulates reproductive fitness in *C. elegans*

**DOI:** 10.1101/2024.12.04.626928

**Authors:** Nicole J. Braun, Danielle E. Mor

## Abstract

The human gut microbiome has attracted considerable attention in recent years due to its broad impact on various physiological processes; however, the mechanisms underlying host-microbiota interactions are not fully understood. In particular, direct causal relationships between specific species and host health outcomes remain to be established. Here we examined the effects of human gut microbiome species *Levilactobacillus brevis* (*L. brevis*) on host health using *C. elegans* and found that *L. brevis* feeding caused significant reproductive dysfunction, including severe egg retention leading to fewer eggs laid, and abnormal egg morphologies. These effects were associated with impaired serotonin signaling in hermaphrodite-specific neurons (HSNs), which regulate egg-laying, while vulval muscle function remained intact. Progeny from *L. brevis*-fed worms exhibited reduced viability and delayed development, suggesting an overall reduction in *C. elegans* reproductive fitness. Our findings align with emerging evidence linking gut dysbiosis to reproductive health disorders in humans, and underscore the need to explore specific roles of individual gut microbiota in host health and physiology. Our work also highlights the utility of *C. elegans* as a model for studying the complex interactions between the microbiome and the host.

## Introduction

The human gut microbiome has gained significant appreciation in recent years for its influence on a wide range of host physiological processes. Beyond its well-known role in digestion, the gut microbiome actively participates in regulating immune, metabolic, and endocrine functions, shaping host physiology across lifespan (1-4). Emerging evidence suggests that disruptions in the gut microbiome—referred to as dysbiosis—are linked to a variety of pathologies, including neurodegenerative diseases, mental health disorders, and reproductive dysfunctions (5-11). Furthermore, gut dysbiosis is increasingly recognized as a contributing factor to reproductive health disorders, including polycystic ovary syndrome (PCOS), endometriosis, and gynecologic cancers (11-12). Recent studies suggest that microbial imbalances in the gut may influence hormonal regulation, immune responses, and metabolic pathways that are critical for maintaining reproductive function (12-13). Dysbiosis may thus serve as a potential underlying factor in the pathogenesis of these disorders, with gut microbiota serving as both a modulator and a potential target for interventions. However, the mechanisms through which the microbiome affects these processes remain poorly understood.

Lactic acid bacteria (L.A.B.) are known inhabitants of the human gut microbiome and are among the bacterial species that undergo diversity changes contributing to gut dysbiosis observed with numerous health conditions (2). L.A.B. are widely used in consumable probiotics and fermented foods under the general assumption that they are uniformly beneficial; however, the lack of current knowledge focusing on specific L.A.B. species leaves a major gap in understanding the interactions of these ubiquitous gut microbiota with the host. One such species, *Levilactobacillus brevis* (*L. brevis*), has been found to be elevated in the gut microbiome of Parkinson’s disease patients (14). In other studies, supplementation with a mix of L.A.B. strains including *L. casei*, *L. acidophilus*, and *L. brevis* caused improvements in mood, behavior, and symptoms of depression in humans, potentially through the production of γ-aminobutyric acid (GABA) (15-16). Understanding how specific microbial shifts, such as those in L.A.B., influence host physiology across various organs and systems is crucial for developing therapeutic strategies aimed at restoring microbiome balance and mitigating disease. Yet, the complexity of the mammalian gut microbiome in tandem with low tractability and slow aging of rodent model systems has made it difficult to establish clear cause-and-effect relationships between specific L.A.B. species and host physiology or disease state.

The small model organism, *C. elegans*, is an excellent system in which to explore host-microbiota interactions given that worms can survive on singular bacterial species allowing for a highly controlled, reductionist approach to determining the causal roles of individual species on host outcomes. In the wild, *C. elegans* maintain a natural, but minimal, gut microbiome consisting of the bacterial species they feed on, including L.A.B. (17). While the *C. elegans* digestive tract is anatomically simple, it offers a streamlined and controlled system with which to uncover species-specific mechanisms of host-microbiota interactions that are not feasible in higher order model organisms. *C. elegans* is also an exceptional model system for the study of host healthspan and offers the unique ability to conduct rapid, mechanistic studies due to having a short lifespan of 2-4 weeks. Moreover, the well-defined spatiotemporal layout of the *C. elegans’* reproductive system combined with the transparency of its tissues make it a powerful system for the study of reproduction and development within the germline (18).

In this study, we show that *C. elegans* raised on *L. brevis* bacteria from the onset of adulthood manifest severe reproductive deficits, including elevated egg retention, an increased proportion of unhatched eggs among those that are laid, and progeny with developmental delay. Furthermore, we find that the serotonergic hermaphrodite-specific neurons (HSNs) of the egg-laying circuitry are both structurally and functionally impaired in *L. brevis*-fed worms, suggesting a critical role of the gut microbiome in regulating neuronal and reproductive health.

## RESULTS

### Feeding *L. brevis* to *C. elegans* increases egg retention in the uterus and causes morphological defects in the retained eggs

Since *C. elegans* are able to feed on singular bacterial species, we took advantage of the ability to establish cause-and-effect relationships between individual gut microbiota and host health. To understand how the human gut microbiome species *L. brevis* impacts *C. elegans* reproduction, non-transgenic worms were first allowed to hatch and develop on the standard laboratory food source of *E. coli* (strain OP50). During the final developmental stage (the L4 stage), the worms were moved to new plates seeded with either *E. coli,* or *L. brevis* derived from a human fecal isolate. This timeline was chosen in order to avoid potential effects on worm development.

On day 4 of adulthood, *L. brevis*-fed worms exhibited a striking increase in egg retention compared to *E. coli*-fed worms (Fig 1A-B). Specifically, *L. brevis*-fed worms retained significantly more eggs inside the uterus than *E. coli*-fed worms with an average of 25 eggs for *L. brevis* compared to 11 eggs for *E. coli*, representing over a 120% increase in egg retention (Fig 1C). Consistent with this result, we observed that over the first four days of adulthood, *L. brevis*-fed worms laid significantly fewer eggs than *E. coli*-fed controls (Supp Fig 1). We next conducted a thorough examination of the morphological characteristics of the eggs retained within the uterus of day 4 *L. brevis*-fed worms. We found that *L. brevis* worsens both the organization and the appearance of the retained eggs, as evidenced by higher incidences of disorganized, irregularly shaped, and abnormally small eggs compared to those of *E. coli*-fed worms (Fig 1D).

**Figure 1.**
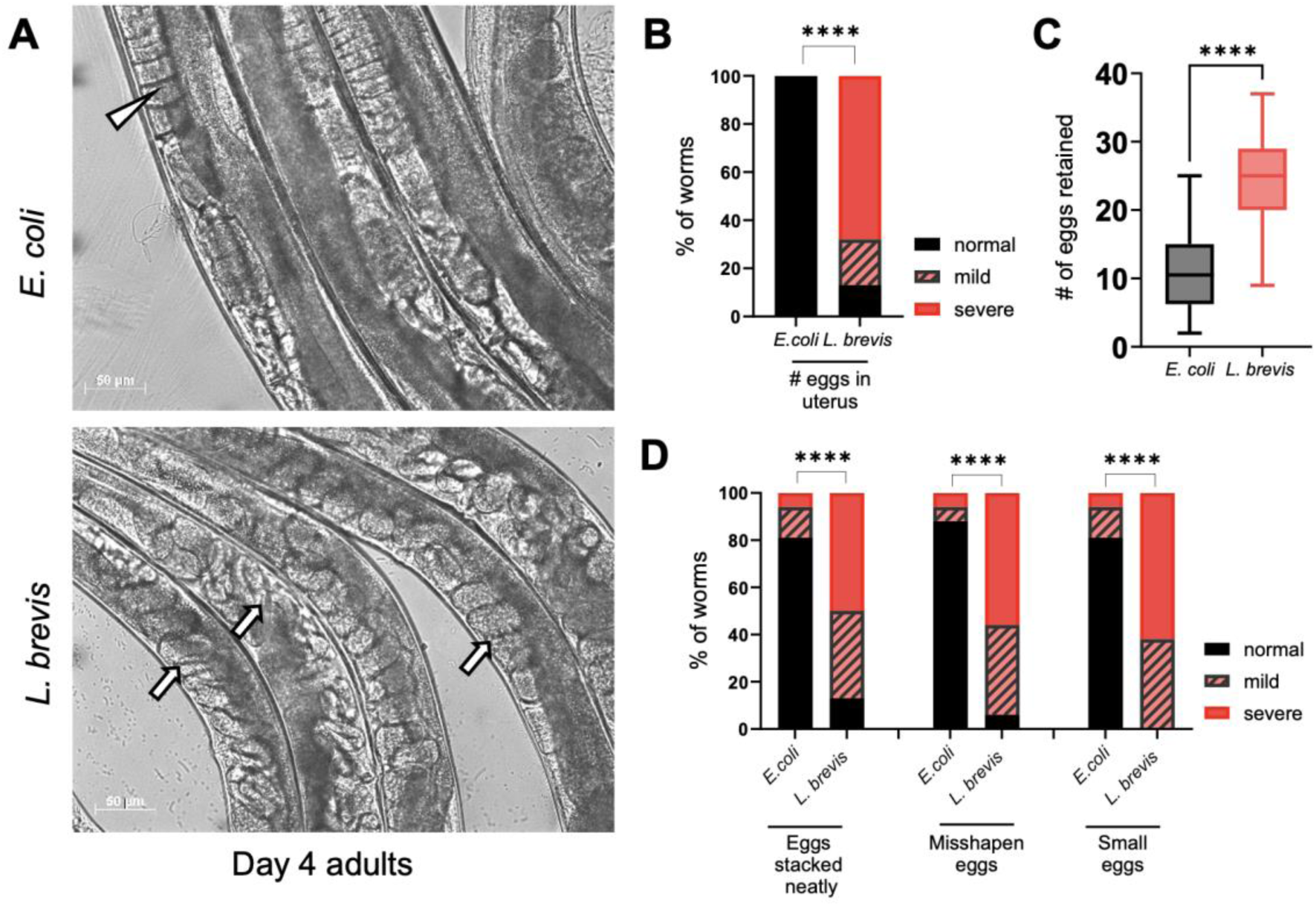
*L. brevis*-fed worms retain their eggs. **(A)** Representative images of eggs in non-transgenic N2 worms fed either *E. coli* (top) or *L. brevis* (bottom); white arrowhead depicts unfertilized oocytes, white arrows depict fertilized embryos (eggs), scale bar = 50µm. **(B)** Scoring of images for normal, mild, or severely abnormal numbers of eggs shows that *L. brevis*-fed worms have severe egg retention (n=16 per group). **(C)** Quantification using the egg-in-worm assay shows that *L. brevis*-fed worms (n=56) have significantly more retained eggs than *E.coli*-fed worms (n=60) on day 4 of adulthood. Unpaired t-test. **(D)** Scored morphologies of the retained eggs show significantly worsened egg organization and appearance in *L. brevis*-fed worms on day 4 of adulthood compared to *E. coli*-fed worms. * p < 0.05; ** p < 0.01; *** p < 0.001; **** p < 0.0001.

### *L. brevis*-fed worms are not starved and have no significant change in lifespan

To validate that the observed reproductive phenotype is not due to starvation in the possible event of *L. brevis* being a non-preferred food source, non-transgenic worms were raised in the absence of food (starved) and directly compared to *L. brevis*-raised worms at day 4 of adulthood (Fig 2). Worms grown on *L. brevis* were found to be morphologically distinct from starved worms (Fig 2A), with starved worms showing significantly lower levels of pigmentation due to the lack of food (Fig 2B). We further confirmed that the worms do indeed ingest *L. brevis* by visually detecting individual rod-shaped *L. brevis* cells stained with acridine orange inside the worm’s buccal cavity (Fig 2C).

**Figure 2.**
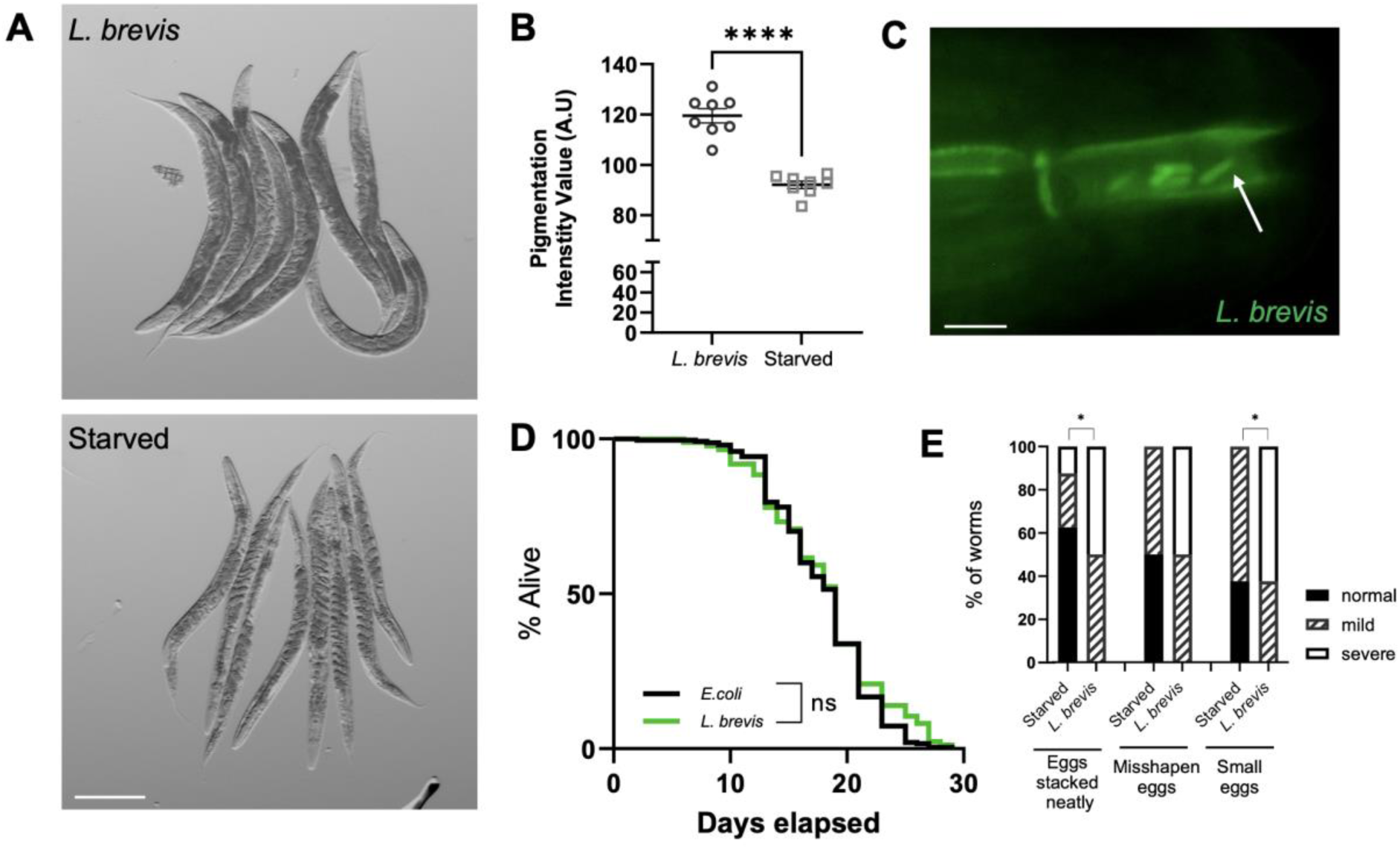
*L. brevis*-fed worms are not starved. **(A)** Day 4 adults raised on *L. brevis* (top) or no food (bottom) from the onset of adulthood. Scale bar = 200µm**. (B)** Quantification of whole-worm pigment intensity. Unpaired t-test, data are mean ± SEM. A.U., arbitrary units. **(C)** *C. elegans* ingest *L. brevis* as visualized with acridine orange; arrow depicts bacterial cells in the mouth of the worm. Scale bar = 5µm. **(D)** No significant difference in lifespan between *E. coli*-fed (n=245) and *L. brevis*-fed (n=96) worms. Log-rank Mantel-Cox survival curve analysis. ns, not significant. **(E)** Scored morphologies of eggs in the uterus show significant differences between *L. brevis*-fed and starved worms. * p < 0.05; ** p < 0.01; *** p < 0.001; **** p < 0.0001.

Additional evidence that worms raised on *L. brevis* do not experience dietary restriction comes from a lack of significant difference in the survival between worms on *L. brevis* versus *E. coli*. Therefore, there is no lifespan extension that is a well-known consequence of starvation or caloric restriction in *C. elegans* (19-22) (Fig 2D). We also found that the abnormal egg morphologies associated with the egg retention phenotype in *L. brevis*-fed worms were not recapitulated by starvation (Fig 2E).

### *L. brevis* causes structural and functional defects in the HSN neurons of the egg-laying circuitry

Egg-laying behavior in *C. elegans* is controlled by a relatively simple circuit consisting of six cholinergic Ventral C neurons (VCs) and two serotonergic Hermaphrodite Specific Neurons (HSNs), which synapse onto the surrounding vulval muscles whose contraction results in the expulsion of eggs (Fig 3A) (23-25). HSN activity and the release of serotonin excites the vulval muscles and the cholinergic VC neurons, with the VC activity directly linked to vulval muscle contractions (23-25). In order to investigate whether the egg retention phenotype induced by *L. brevis* may be caused by defects in specific components of the egg-laying circuit, worms were exposed to serotonin or levamisole, which are both pharmacological agents previously shown to stimulate egg-laying (26-27). On day 4 of adulthood, *L. brevis*-fed worms exposed to exogenous serotonin were able to lay eggs with no significant difference from the *E. coli*-fed controls (Fig 3B). This indicated that the vulval muscle itself is intact and able to respond to serotonin with contraction leading to successful egg-laying. Since the application of exogenous serotonin bypasses the activity of the HSN, these data suggest that a lack of endogenous serotonin signaling from the HSNs may be responsible for the egg retention phenotype.

**Figure 3.**
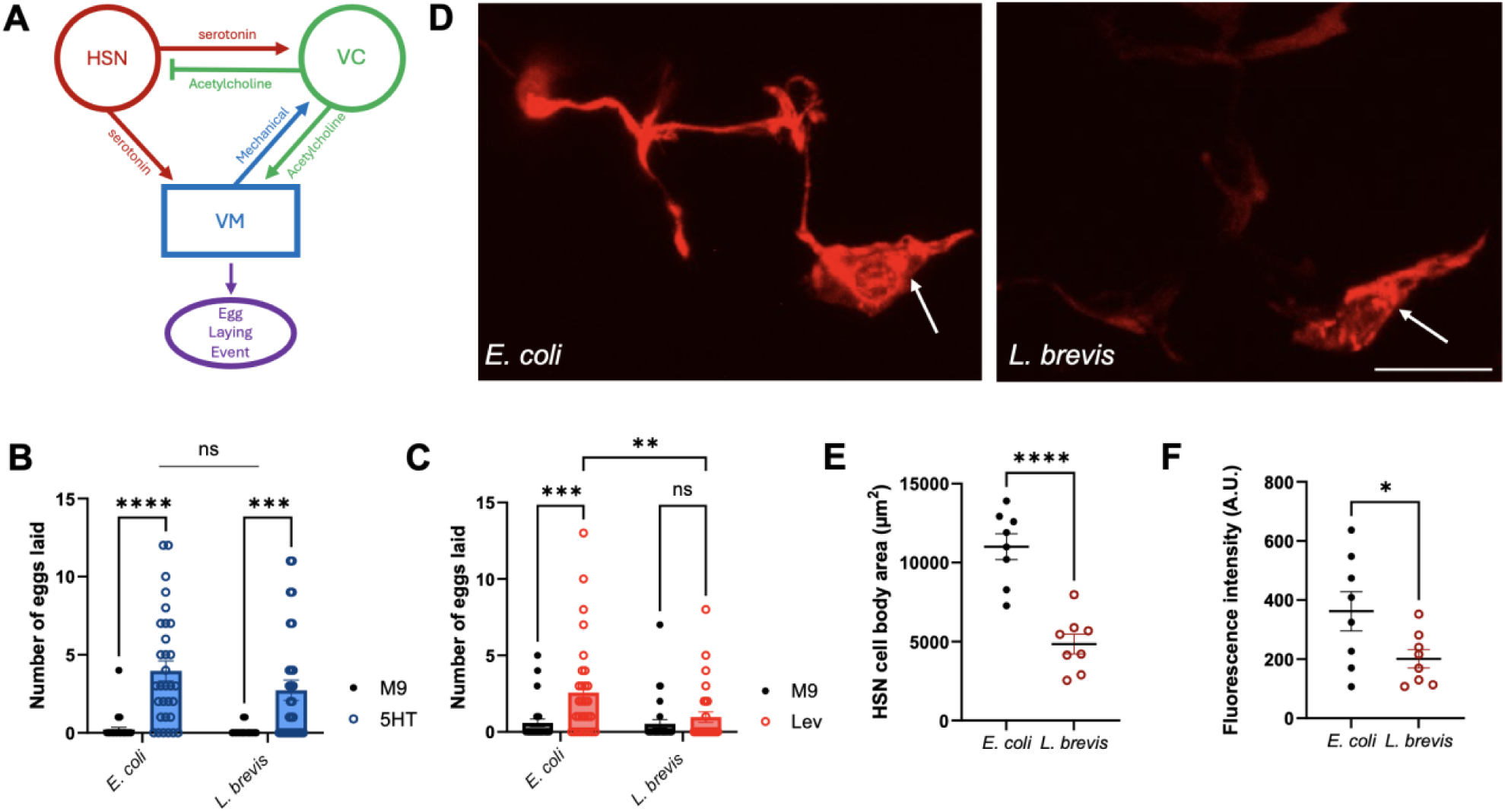
HSN structural and functional defects in worms fed *L. brevis*. **(A)** Simplified schematic of the *C. elegans* egg-laying circuitry. HSN, hermaphrodite-specific neurons. VC, Ventral C neurons. VM, vulval muscle. **(B)** Eggs laid in response to exogenous serotonin (5HT) or M9 treatment of day 4 adults (n=30 per group). Two-way ANOVA, data are mean ± SEM. ns, not significant. **(C)** Eggs laid in response to levamisole or M9 treatment of day 4 adults (n=30 per group). Two-way ANOVA, data are mean ± SEM. ns, not significant. **(D)** Representative images of the HSN in day 4 adults raised on *E. coli* (left, n=10) or *L. brevis* (right, n=10), with arrow depicting the cell body. Scale bar = 50µm**. (E-F)** Quantification of HSN cell body area **(E)** and fluorescence intensity **(F)**. Unpaired t-tests, data are mean ± SEM. A.U., arbitrary units. * p < 0.05; ** p < 0.01; *** p < 0.001; **** p < 0.0001.

To further test the function of the HSNs, we used the acetylcholine receptor agonist, levamisole, to stimulate egg-laying and found that in this case *L. brevis*-fed worms did not show enhanced egg-laying, in contrast to *E. coli*-fed worms (Fig 3C). Previous studies have suggested that serotonin from the HSN is essential to enhance VC and vulval muscle activity and to coordinate the process of egg-laying. In the absence of HSN-potentiation, the VC neurons alone are unable to stimulate the vulval muscles to the threshold required for laying eggs. Since the application of exogenous levamisole effectively bypasses VC activity, the lack of enhanced egg-laying further suggests that HSN neuronal activity is impaired.

Together our findings indicate that *L. brevis* impacts the function of serotonin signaling by the HSNs, likely contributing to the egg retention phenotype observed in *L. brevis*-fed worms. This was further corroborated with fluorescence imaging of the HSNs, which revealed structural defects in the *L. brevis*-fed worms compared to *E. coli* controls at day 4 of adulthood (Fig 3D). Examination of the HSNs showed significantly smaller cell bodies in *L. brevis*-fed worms compared to *E. coli* controls, in conjunction with decreased HSN fluorescence intensity (Fig 3E-F).

### *L. brevis*-fed worms have significantly fewer live progeny, reduced embryonic viability, and progeny have developmental delays

To assess the fitness of the progeny of hermaphrodite ‘mothers’ raised on *L. brevis,* we performed viability assays throughout the first four days of adulthood when *C. elegans* are most reproductively active (Fig 4A-D). Across days 1-4, the total number of live progeny per parent worm was significantly reduced in *L. brevis*-fed worms compared with *E. coli*-fed controls (Fig 4A). Throughout the experimental period, *L. brevis*-fed worms produced significantly fewer live (hatched) progeny on each day from days 1-3 (Fig 4B), and as well as significantly greater unhatched embryos (Fig 4C) on each day from days 1-4, compared to *E. coli*-fed controls. Embryonic viability was calculated as the number of live progeny divided by the total progeny (live and unhatched) across the experimental period. *E. coli*-fed controls yielded an embryonic viability of 98% while *L. brevis*-fed worms showed a substantial reduction to only 63% (Fig 4D). These findings show that, in combination with the lower total number of eggs laid by *L. brevis*-fed worms (Supp Fig 1), overall *L. brevis* causes worms to produce fewer live progeny, and for a greater proportion of the eggs they lay to be unable to hatch.

**Figure 4.**
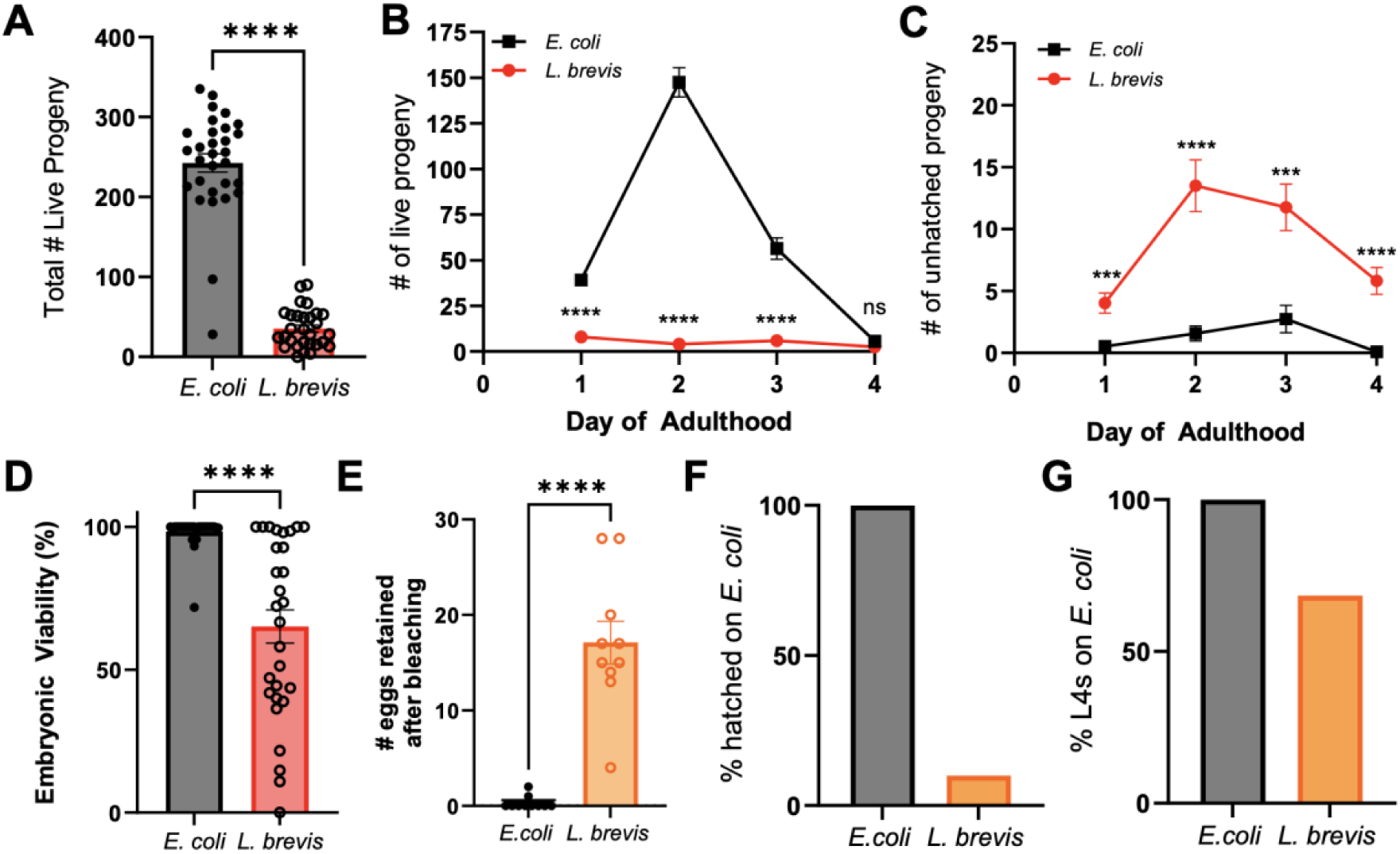
Progeny of worms raised on *L. brevis* have hatching and developmental defects. **(A)** Total number of live progeny produced from the first 4 days of adulthood is significantly reduced in *L. brevis*-fed worms (n=29) compared with *E. coli* controls (n=30). Unpaired t-test, data are mean ± SEM. **(B-C)** Progeny viability assay over the first 4 days of adulthood, revealing that *L. brevis*-fed worms (n=29) have significantly fewer live progeny produced each day **(B),** and significantly more unhatched embryos produced each day **(C)** than *E. coli* controls (n=30). **(D)** Embryonic viability was substantially decreased in the *L. brevis*-fed population (n=29) compared to *E. coli* (n=30) on day 4 of adulthood. Unpaired t-test, data are mean ± SEM. **(E)** Number of eggs retained after picking day 4 worms into a drop of bleach on *E. coli*-seeded plates (n=10 per group). Unpaired t-test, data are mean ± SEM. **(F-G)** Percent of eggs hatched **(F)** and developed to L4 stage **(G)** after bleaching day 4 *L. brevis-* or *E. coli*-fed worms onto *E. coli*-seeded plates. * p < 0.05; ** p < 0.01; *** p < 0.001; **** p < 0.0001

Finally, we asked if the decline in progeny viability was due to the eggs being laid in the presence of *L. brevis,* possibly contributing to an unfavorable environment for hatching. To address this, *C. elegans* were raised as described previously to day 4 of adulthood before being picked into a drop of hypochlorite solution on an *E. coli*-seeded plate to “collect” the retained eggs. This allowed the eggs to hatch in a known favored environment. After bleaching, *L. brevis*-fed worms retained 171 eggs compared to only 3 from *E. coli*-fed worms that were left to hatch on the *E. coli*-seeded plates (Fig 4E). 24 hours later, 100% of the eggs from parents raised on *E. coli* had hatched, while only 11% of the eggs from parents raised on *L. brevis* had hatched (Fig 4F), suggesting that the *L. brevis*-induced decline in progeny viability is due to intrinsic factors of the eggs rather than their environment. We further assessed the developmental rate of the progeny that had hatched on *E. coli*-seeded plates. At 20°C, synchronized *C. elegans* will develop to the L4 larval stage within approximately 48 hours. At 48 hours after transferring the eggs to the *E. coli*-seeded plates by bleaching, 100% of the progeny from parents raised on *E. coli* developed to the L4 stage while only 68% of the *L. brevis* progeny developed to L4, indicating a developmental delay.

## Discussion

While the gut microbiome consists of a diverse community of over 1,500 bacterial species that contribute to nutrient metabolism, pathogen defense, and intestinal barrier integrity (1), its complex interactions with distant organs and systems—such as the immune, cardiovascular, and reproductive systems (2-3, 28-29) — suggest that its influence extends far beyond digestion. These microbial communities also engage in bidirectional communication with the host through neuronal, endocrine, and immune signaling pathways (29-30). Such interactions underscore the microbiome’s essential role in maintaining systemic homeostasis. Our study provides new insights into the effects of *L. brevis* on host health, particularly focusing on reproduction in *C. elegans*. Our findings highlight that feeding *L. brevis* to *C. elegans* leads to significant alterations in reproductive outcomes, including increased egg retention, reduced egg-laying, and developmental defects in both embryos and progeny. These observations suggest that specific gut microbiota species can modulate host physiology in ways that may have implications for understanding reproductive health in humans, particularly in the context of gut dysbiosis. Given the increasing recognition of gut microbiota dysbiosis as a contributing factor in reproductive pathologies in humans, including PCOS, endometriosis, and chronic anovulation (11-12, 31), our findings support the growing notion that microbial shifts in the gut—whether due to dietary, environmental, or genetic factors—could have profound effects on reproductive health.

Our finding that *L. brevis* reduced reproductive fitness in *C. elegans* aligns with a previous study that observed low reproductive and developmental rates in *C. elegans* fed 35 different strains of L.A.B. (32). Several studies across different model systems support the idea that L.A.B. can influence various aspects of reproduction. For instance, L.A.B. have been shown to alleviate symptoms of polycystic ovary syndrome (PCOS) in rat models by modulating the gut microbiota through the regulation of sex hormones (33). Similarly, a study in zebrafish demonstrated that feeding *L. casei* and *L. rhamnosus* improved both fecundity and immunity. However, these beneficial effects were lost once the probiotics were removed from the diet (34). Additionally, research in Drosophila highlighted both direct and trans-generational effects of the gut microbiota on reproduction (35). Furthermore, germ-free mice showed an increase in reproductive capacity after exposure to bacteria (36), reinforcing the idea that the gut microbiota plays a crucial role in regulating reproductive health. These studies together with our findings underscore the need to better understand the specific effects of individual bacterial species, including *L. brevis*, on host physiology.

In conclusion, our study provides compelling evidence that *L. brevis* can disrupt host reproductive function in *C. elegans*, potentially through alterations in serotonin signaling and HSN neuron function. These findings lay the groundwork for further studies exploring the role of specific microbiota species in regulating reproductive health and the broader implications of gut dysbiosis for human disease. Given the growing evidence of the gut microbiome’s influence on both local and distal bodily functions, restoring microbial balance could emerge as a promising avenue for mitigating disease and promoting overall health across the lifespan.

## Materials and Methods

### *C. elegans* strains and maintenance

*C. elegans* strains were maintained at 20°C under standard conditions (34). Hypochlorite alkaline-bleaching solution was used to synchronize worms for experiments (34). Briefly, synchronization was performed by exposing gravid hermaphrodites to the hypochlorite solution to collect eggs, followed by thorough washing of the collected eggs in M9 buffer and dispensing onto high growth medium (HG) plates seeded with OP50 *E. coli*. For experiments, standard nematode growth medium (NGM) plates were seeded with 1 mL of either *E. coli* or *L. brevis* for *ad libitum* feeding. The following strains were used: wild-type N2 Bristol strain, and LX975 vsls13 [*lin-11::pes-10::GFP + lin-15(+)*]; vsIs97 [*tph-1p::DsRed2 + lin-15(+)*]; vsIs100 [*myo-3p::CFP + lin-15(+)*].

### Bacterial growth conditions

The *L. brevis* strain used in experiments was obtained from Microbiologics (Catalog No. 01262L) originally derived from ATCC 14869 isolated from human fecal samples. For these experiments, *L. brevis* was cultivated in deMan, Rogosa, and Sharpe (MRS) medium at 35°C for 24-48 hours and stored at 4°C. OP50 *E. coli* was grown at room temperature overnight as per standard protocols (34).

### Egg-in-worm assay

Day 4 adult hermaphrodites were placed individually into 10μL of hypochlorite solution and eggs were counted after the body of the worm dissolved (roughly 10 minutes). Protocol was followed as described previously (38).

### Scoring egg quality

Worms were mounted on 2% agarose pads with M9 and sodium azide and imaged on a Leica M165 FC fluorescent stereomicroscope with a Leica K5 microscope camera. Brightfield images were used to characterize the morphology of eggs retained within the uterus. Using a similar method to previous characterization of oocyte morphology (39), images were scored based on the following criteria: a score of normal, mild or severe was assigned for each category based on the severity of the phenotype. In regard to the ‘eggs in uterus’ category, normal indicated 0-15 eggs in varying developmental stages, mild indicated 16-20 eggs with some fully developed embryos, and severe indicated greater than 20 eggs retained with nearly all fully developed or the presence of matricidal hatching; with respect to the ‘eggs stacked neatly’ category, normal indicated eggs, if present, were organized or “stacked” neatly inside the uterus, mild indicated some slight irregularities where some eggs appear overlapped, and severe indicated the eggs appear disorganized and crowded; with respect to the ‘misshapen eggs’ category, normal indicated no abnormally shaped eggs, mild indicated the presence of slight irregularities, and severe indicated most eggs appeared irregularly shaped or damaged; with respect to the ‘small eggs’ category, normal indicated no small eggs, mild indicated some eggs were smaller than others, and severe indicated that most of the eggs appear very small.

### DIC microscopy

Worms were mounted on 2% agarose pads with M9 and sodium azide and imaged on a Zeiss Axioplan 2 Fluorescence Microscope with differential interference contrast (DIC) at 10x magnification.

### Progeny viability assays

This assay was used to track egg-laying and progeny-hatching in adult worms over the first four days of adulthood. On Day 0, synchronized L4 stage worms were individually plated onto 35mm NGM plates seeded with either *E. coli* or *L. brevis*. The assay involved moving individually plated worms to new plates daily and counting the number of eggs laid, live hatched progeny, and unhatched embryos until the initial worm reached day 4 of adulthood. Worms were censored due to matricide (progeny hatched inside the hermaphrodite parent), vulval rupture (intestinal expulsion), or if lost. Censored animals were not included in the data. The average number of live progeny per worm was calculated by summing the number of progeny produced and dividing by the number of parent worms. Embryonic viability was calculated as the number of live progeny divided by the total progeny (live and unhatched) across the experimental period. Modified from (39).

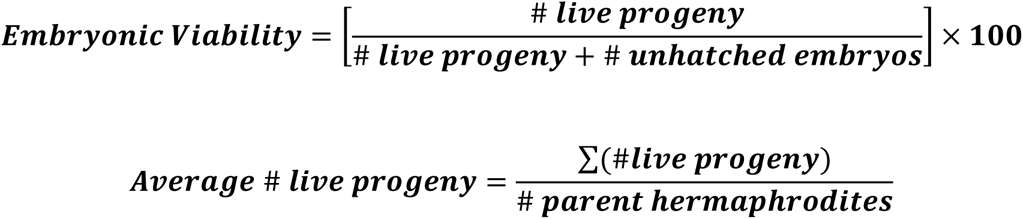

### Survival assay

For lifespan assays, as previously outlined (40), groups of 12-15 L4 stage hypochlorite-synchronized worms were initially placed on individual NGM plates, seeded with either *E. coli* or *L. brevis*. Worms were transferred to freshly seeded plates every two days while producing progeny, and every 3 days thereafter. Worms were considered dead when they no longer responded to touch stimuli and were censored in the event of matricide (progeny hatched inside the hermaphrodite parent), vulval rupture (intestinal expulsion), or if lost. Censored animals were not included in the data.

### Acridine orange bacteria staining

*L. brevis* cells were collected into pellets by centrifugation (4000g x 10 mins). The pellet was washed in PBS three times before being resuspended in acridine orange for 15 minutes. 500μL of the acridine orange-stained bacteria was seeded on an NGM plate and allowed to dry before worms were placed into the bacteria. After one hour, the worms were mounted on 2% agarose pads with M9 and sodium azide and imaged at 40x magnification with a GFP filter.

### HSN neuron imaging

Worms expressing dsRED in HSN neurons (strain LX975) were mounted on 2% agarose pads with M9 and sodium azide and imaged on a Nikon A1R MP+ multiphoton/confocal microscope at 60x magnification. Maximum intensity projections were generated, and a region of interest was drawn around each cell body to measure cell body area (μm2) and fluorescence intensity. LUTS settings were standardized across images.

### Pharmacologically induced egg-laying

Day 4 adult hermaphrodites were placed in individual microtiter wells containing 50μL of a solution of either serotonin (5mg/mL) or levamisole (0.1mg/mL) dissolved in M9 buffer and the number of eggs released in each well were counted after 60 mins. 10 worms were tested per drug per experiment.

### Statistical analysis

Statistical analyses were performed using GraphPad Prism 10.2. An unpaired two-tailed Student’s t-test was used for all comparisons between two groups. For comparisons between multiple groups, One-Way ANOVA or Two-Way ANOVA (for repeated measures or two-variable analyses) was performed with post-hoc testing as indicated.

## Acknowledgments

We thank the *Caenorhabditis* Genetics Center for the LX975 strain (P40 OD010440). The N2 strain was generously provided by C. Murphy (Princeton University). We thank the Department of Neuroscience and Regenerative Medicine 2-Photon Microscopy Facility, and the Medical College of Georgia Imaging Core Facility. This work was supported by the start-up fund from the Medical College of Georgia at Augusta University.

## Author Contributions

N.J.B. and D.E.M. conceived and designed the experiments. N.J.B. performed the experiments and analyzed the data. N.J.B. and D.E.M. wrote the paper.

## Declaration of Interests

The authors declare no competing interests.

## Notes

### Competing Interest Statement

The authors have declared no competing interest.

