## Supplemental Figure 1 for "Gut microbiome species *Levilactobacillus brevis* regulates reproductive fitness in *C. elegans*"

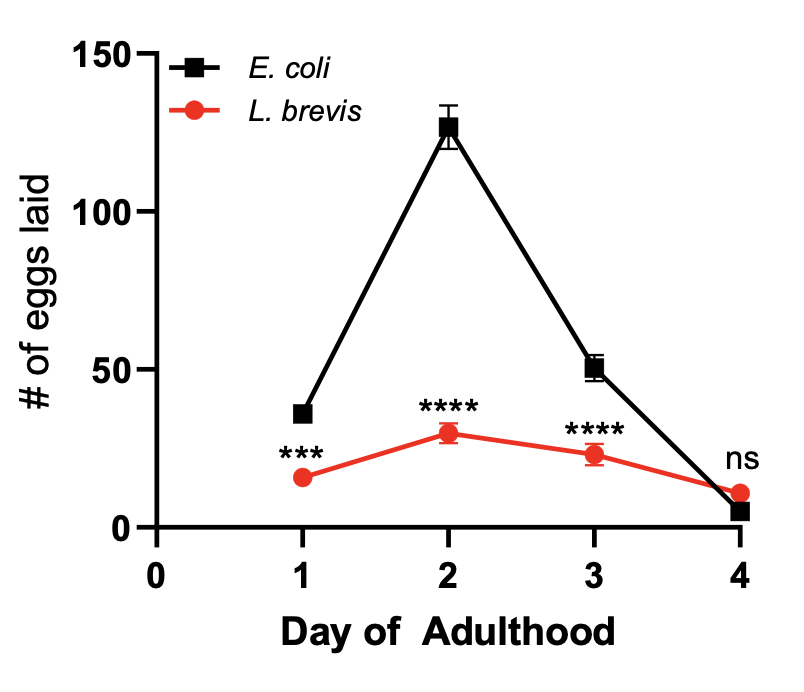

**Supplemental Figure 1. *L. brevis*-fed worms lay fewer eggs.** Progeny viability assay over the first 4 days of adulthood revealed that *L. brevis*-fed worms (n=29) laid significantly fewer eggs than *E. coli*-fed worms (n=30). ns, not significant. * p < 0.05; ** p < 0.01; *** p < 0.001; ****p < 0.0001
